# Endothelial Sensitivity to Pro-Fibrotic Signals Links Systemic exposure to Pulmonary Fibrosis

**DOI:** 10.1101/2025.01.02.631148

**Authors:** Lena Möbus, Laura Ylä-Outinen, Luca Mannino, Giorgia Migliaccio, Karoliina Kosunen, Nicoletta D’Alessandro, Angela Serra, Dario Greco

**Affiliations:** Finnish Hub for Development and Validation of Integrated Approaches (FHAIVE), Faculty of Medicine and Health Technology, Tampere University, 33520 Tampere, Finland; Tampere Institute for Advanced Study, Tampere University, 33520 Tampere, Finland; Division of Pharmaceutical Biosciences, Faculty of Pharmacy, University of Helsinki, 00790 Helsinki, Finland

**Keywords:** Endothelial cells, pulmonary fibrosis, adverse outcome pathway, toxicology, pro-fibrotic substances, mechanism of action, transcriptomics, toxicogenomics, *in vitro*

## Abstract

Pulmonary fibrosis (PF) is a life-threatening condition characterised by excessive extracellular matrix deposition and tissue scarring. While much of PF research has focused on alveolar epithelial cells and fibroblasts, endothelial cells have emerged as active contributors to the disease initiation, especially in the context of systemic exposure to pro-fibrotic substances. Here, we investigate early transcriptomic and secretory responses of human umbilical vein endothelial cells (HUVEC) to subtoxic doses of bleomycin, a known pro-fibrotic agent, and TGF-beta, a key cytokine in fibrosis. Bleomycin exposure induced a rapid and extensive shift in the endothelial transcriptional programme, including signatures of endothelial to mesenchymal transition, cellular senescence, and immune cell recruitment. These findings suggest endothelial cells as early initiators of pro-fibrotic signals, independent of contributions from other cell types. In contrast, TGF-beta effects were limited and transient, indicating its pro-fibrotic action may require another initial stimulus and interplay with other cells like fibroblasts. This study highlights the sensitivity of endothelial cells to systemic pro-fibrotic exposure and provides a blueprint of early pro-fibrotic mechanisms, emphasising their pivotal role in PF pathogenesis.

## Introduction

Fibrosis, causing up to 45% of deaths due to organ failure, involves excessive collagen and extracellular matrix (ECM) deposition, leading to tissue scarring and finally loss of tissue function.^1–5^ Pulmonary fibrosis (PF) is notably prevalent and fatal, affecting 10-60 per 100,000 overall and up to 400 per 100,000 in those over 65. It is driven by factors like air pollution and certain drugs but also unknown factors.^6^ PF can be initiated externally by inhaled stimuli affecting alveolar epithelial cells and triggering immune responses and fibroblast activation, or internally via the pulmonary endothelium, which then orchestrates downstream processes in the lung interstitium. In the past, pro-fibrotic stimuli exerting their effects from the outside compartment of the lungs have been more extensively studied with especially fine particulate matter, carbon nanotubes, asbestos and titanium dioxide being known inhalation hazards.^7,8^ More recent research is stressing the crucial and active role of endothelium in various diseases, and the awareness that endothelial cells are not only responsible for forming the barrier that compartmentalises blood from tissues but exert an active role in metabolism and are drivers of physiological and pathophysiological processes.^9^

One of the best studied pro-fibrotic substances is bleomycin, a cytostatic glycopeptide whose chemical properties ultimately result in DNA breaks. Used as a chemotherapeutic for several malignancies, it is associated with drug-induced interstitial pneumonitis and PF as spontaneous adverse outcome.^10^ Bleomycin is further used to induce experimental PF in rodents by tracheal administration of the chemical.^11,12^ Importantly, in the clinical context, bleomycin is administered systemically. Thus, there is evidence that bleomycin exerts pro-fibrotic effects via both the external (alveolar epithelium) and the internal (alveolar endothelium) entrance to the lungs. When administered systemically, the alveolar microvasculature by design is substantially exposed to the chemical given that the lungs receive 100% of the cardiac output within short time. The role of endothelial cells as mediators in fibrosis is now acknowledged. Endothelial cellular processes observed in the context of fibrosis in clinical and animal studies have been summarised under endothelial to mesenchymal transition (EndMT), cellular senescence, fibroblast activation and immune cell recruitment by secretion of pro-fibrotic and pro-inflammatory mediators, as well as vascular rarefaction and capillary leaking.^13,14^

Several candidate biomarker studies investigated the effects of pro-fibrotic substances directly administered to endothelial cells showing an induction of immune cell recruitment machinery including interleukin-8, E-selectin, and ICAM1.^15–17^ Besides bleomycin, TGF-beta, a key driver of tissue repair that becomes dysregulated in fibrosis, has been used to model processes such as EndMT as demonstrated by upregulation of SNAIL and ACTA2 and downregulation of CD31 and tight junction machinery.^18–21^ Aside from a few *in vitro* candidate biomarker studies, most investigations of endothelial cells in the context of fibrosis have been conducted *in vivo* focusing on advanced stages of PF. Of those, most studies used a rodent model with intratracheal instillation of bleomycin or TGF-beta triggering externally initiated PF leaving endothelial cells rather as secondary reactors to stimuli by other cell types that had direct contact with the pro-fibrotic substances.^20,22–27^ Though different routes of bleomycin administration cause PF, mechanistically they are different. One dose of intratracheal administration in rodents is enough to cause PF-like conditions after 14-28 days, which however are self-limiting and resolve after 6 weeks.^11^ With intravenous administration of bleomycin, it takes longer time to establish fibrotic conditions in the lung including alveolar epithelial hyperplasia and therefore it is less practical for *in vivo* studies, although mechanistically it would resemble bleomycin-induced development of PF in humans. It has been hypothesised that bleomycin induces PF in two stages, an early inflammatory and later fibrotic stages.^28^ However, in *in vivo* studies, molecular signatures are derived mostly from already established states of PF, which neglects to study early initiating events.

To characterise immediate mechanisms of endothelial cells exposed to pro-fibrotic substances, we profiled the transcriptome of human umbilical vein endothelial cells (HUVEC) exposed to bleomycin as exogenous as well as TGF-beta as endogenous pro-fibrotic substance, over 3 days and at 3 subtoxic doses. Transcriptome analysis was accompanied by the analysis of secreted factors. We demonstrate that endothelial cells are extremely sensitive to bleomycin, showing a significant shift of the transcriptome involving known PF-related fingerprints such as EndMT and immune cell recruitment signatures. In contrast, TGF-beta had limited effects to the cells both in terms of time and extent, suggesting that TGF-beta alone is not a potent pro-fibrotic stimulus on endothelial cells. Taken together, our results point to a possible starting point of pro-fibrotic mechanisms initiated by the endothelium under systemic exposure regime.

## Results & Discussion

To investigate the early response of endothelial cells to pro-fibrotic stimuli, we performed transcriptome analysis of HUVEC cells grown in a monolayer and exposed to multiple concentrations of bleomycin (0, 7, 14, and 21 µg/ml) as exogenous pro-fibrotic substance, and TGF-beta (0, 5, 10, and 20 ng/ml), known as endogenous pro-fibrotic compound, respectively. We carried out our analysis at 3 consecutive time points by taking samples at 24, 48 and 72 hours of exposure in replicates of 4 for each experimental condition. In addition, we measured the secretion of 22 candidate cytokines and peptidases in response to different doses of bleomycin and TGF-beta and at multiple timepoints to evaluate secreted factors related to the initiation of pro-fibrotic conditions.

### Significant difference between exogenous and endogenous pro-fibrotic stimuli on endothelial cells

Transcriptional data exploration by principal component analysis showed clear dose-dependent (first component) effects of bleomycin exposure, but not for TGF-beta exposure (Figures S1A and S1B). After confirming that both datasets, bleomycin and TGF-beta experiments, had overall non-different numbers of raw counts per sample reflecting the effective sequencing dataset size (Figure 1A), we investigated the distribution of measured expression height over all genes in the unexposed and exposed cells of both datasets. In the unexposed cells, the distribution of expression levels resembled each other in both datasets, as it would be expected (Figure 1B). In contrast, in the exposed cells, we observed different distributions for the bleomycin and TGF-beta exposure (Figure 1C). In bleomycin-exposed cells, a higher number of genes was expressed at low level indicated by low RNA-Seq counts, whereas in TGF-beta-exposed cells, a higher number of genes was expressed at high level. Further, the number of detected genes (>= 1 count in the dataset) differed between the two datasets, with 52,921 and 37,989 detected genes in the bleomycin and TGF-beta dataset, respectively, while the number of genes detected in controls of both datasets was 30,881 and 31,638, respectively. This indicates that bleomycin induces the expression of multiple genes, while under TGF-beta, the overall gene expression profile remains comparable to unexposed cells. In bleomycin-exposed cells, 152 genes were expressed (>=10 counts in at least 4 samples), that were neither detected in unexposed nor TGF-beta-exposed cells (0 counts in all samples) (Table S1). Among those genes exclusively induced by this exogenous pro-fibrotic substance were genes involved in immune-cell chemotaxis (*CCL22* and *CCR3*), tissue remodelling (*ADAMTS20* and *MMP12*), and cell-cell adhesion (*DSG1* and *DSG3*). Upon TGF-beta treatment, no genes were expressed (>=10 counts in at least 4 samples) that were not also detected in the unexposed cells, confirming that TGF-beta does not alter the number of expressed genes as compared to the unexposed cells.

**Figure 1:**
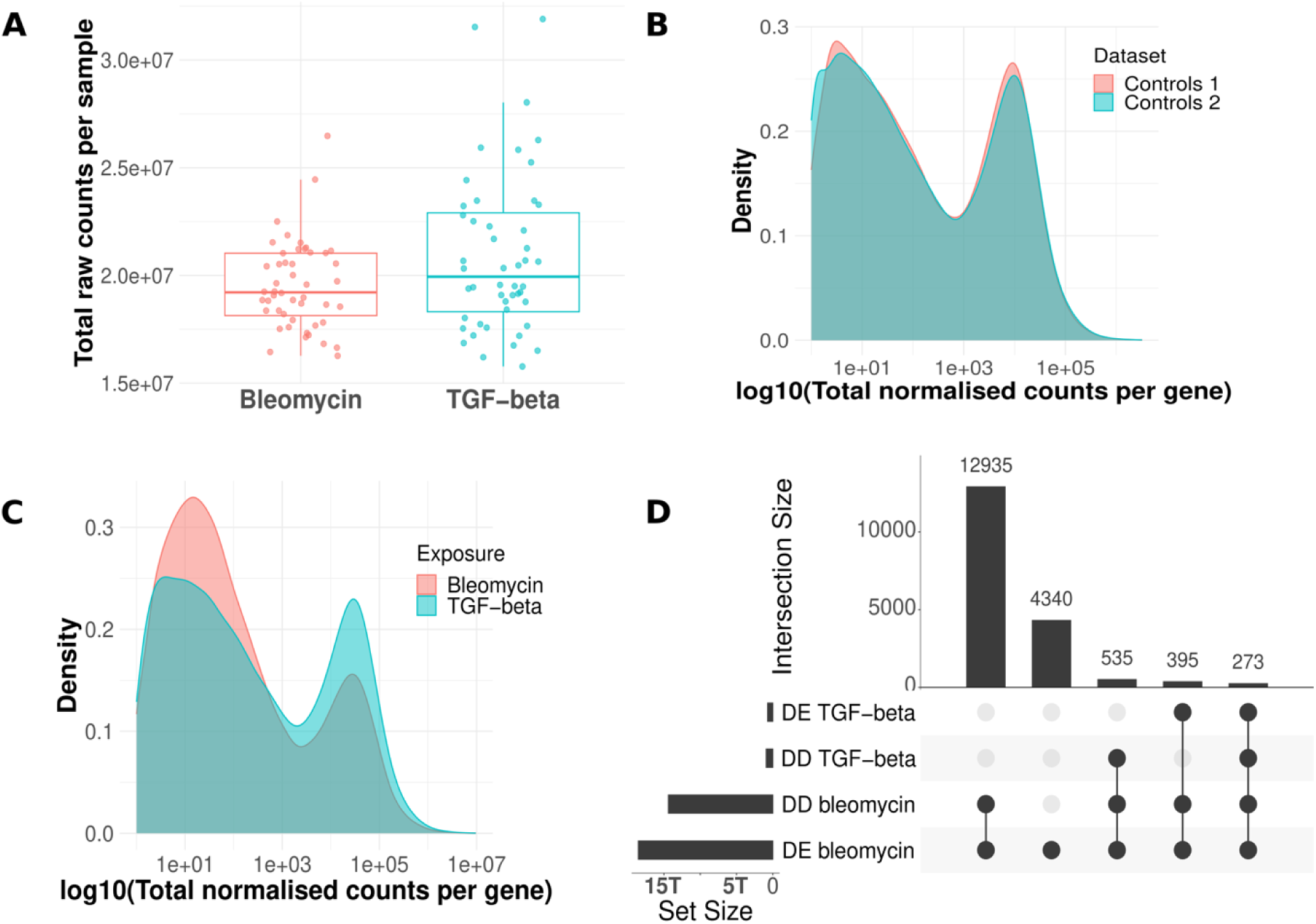
Overview of RNA-Seq data. **A**. Total raw counts obtained per sample for both datasets. Wilcoxon Rank Sum test showed no significant difference in raw counts between the two datasets (p value = 0.12). **B**. Distribution of total normalised counts per gene summed up over all samples (n=12) of unexposed cells in both datasets. **C**. Distribution of total normalised counts per gene summed up over all samples (n=36) of exposed cells in both datasets. **D**. Set size and intersection size of dose-dependent (DD) and differentially expressed (DE) genes in both exposures. The union of DD and DE genes detected in the different experimental sub-groups (timepoints, concentrations) is plotted.

Bleomycin significantly altered the transcription of 35.5 % of all detected genes (18,780 of 52,921 genes, significant by differential expression and/or dose-dependent analysis), while TGF-beta affected only 3.6 % (1,364 of 37,989) of the transcriptome, and the fold changes were overall smaller than for bleomycin (Figures 1D and S2, Table S2-S4). Importantly, the TGF-beta signalling machinery including genes encoding the TGF-beta receptors and key downstream mediators such as SMAD family members was expressed in HUVEC cells (Figure S8). Furthermore, according to public expression databases such as the Human Protein Atlas and Expression Atlas, endothelial cells generally express the TGF-beta receptor-encoding genes, demonstrating their capability to respond to TGF-beta.^29,30^ Indeed, among the overall mild expression changes under TGF-beta, we identified a comparably small set of 39 genes that were dose-dependently altered over all 3 timepoints of the experiment. Sixteen of these genes were either direct interaction partners of *TGFB1* or linked to it via one other partner in the human interactome indicating that TGF-beta causes a narrow persistent response in HUVEC cells centred around TGF-beta signalling, while most other transcriptional changes were of short duration only (Figure S3). The stark difference in transcriptional responses between bleomycin and TGF-beta may be attributed to the nature of the two compounds. TGF-beta is an endogenous signalling molecule that endothelial cells encounter in homeostatic conditions, where their responses are tightly regulated. In contrast, bleomycin is an exogenous cytotoxic agent. Additionally, TGF-beta is primarily known as a key activator of fibroblasts, which in turn can influence endothelial cells through paracrine signalling or matrix remodelling. In our experimental setup, the absence of fibroblasts may render the endothelial response to TGF-beta inherently self-limited. This self-regulation seems particularly relevant at the apical surface of endothelial cells, where TGF-beta was applied in our experiment. *In vivo*, TGF-beta plays a critical role in mediating processes like platelet adhesion and leukocyte recruitment during vascular injury.^31,32^ However, these functions must be carefully modulated and promptly deactivated to prevent excessive inflammation and collateral damage once the injury is resolved. This necessity for transient and restrained activity may explain the limited transcriptional changes observed in endothelial cells exposed to TGF-beta in our study. In another study, TGF-beta exhibited pro-fibrotic effects on endothelial cells only when the cells were derived from fibrotic lung tissue, but not when they originated from healthy lung tissue suggesting that endothelial cells respond to TGF-beta predominantly under pathophysiological conditions, likely due to prior reprogramming associated with the diseased state.^33^

To understand what the responses of HUVEC cells to exogenous and endogenous pro-fibrotic substances have in common, we performed functional enrichment analysis of differentially expressed genes dysregulated in the same direction under both substances. Since there were only 3 dysregulated genes in common at 48 hours, enrichment analysis was performed for 24 and 72 hours only. At 24 hours, commonly upregulated genes upon bleomycin and TGF-beta exposure were enriched for chemokine activity, while downregulated genes were enriched for the process of branching involved in blood vessel morphogenesis indicating an early inflammatory signal together with compromised cellular development (Figure S4, Table S5). At 72 hours, commonly upregulated genes were enriched for ECM related processes, functions, and compartments suggesting remodelling processes of ECM under both substances, whereas downregulated genes were enriched for processes and functions related to cell cycle homeostasis.

Given the remarkably high number of genes affected by an exogenous pro-fibrotic substance, we compared this with our recent observations (Morikka et al., 2024), where we investigated the transcriptomic effects of bleomycin on macrophages on a model of differentiated THP-1 cells. THP-1 macrophages exposed to bleomycin showed 918, 1,119, 1,707, 2,499 and 3,072 differentially expressed genes (adjusted p-value <=0.01, log2 fold change (LFC) >=0.58) at doses of 20, 40, 60, 80, and 100 µg/ml bleomycin after 24h, respectively, whereas the HUVEC cells showed 6,967, 10,608, and 12,741 differentially expressed genes at doses of 7, 14, and 21 µg/ml bleomycin (adjusted p-value <=0.01, LFC >=0.58), respectively.^34^ The doses in both experiments were chosen based on cell viability (IC5 to IC20). Together we conclude that (1) HUVEC cells have a lower threshold for subtoxic doses of bleomycin as compared to THP-1 macrophages, and (2) despite comparable viability measured under the exposure, HUVEC cells present a higher degree of dysregulation in gene expression as compared to THP-1 cells suggesting that endothelial cells are more sensitive to exogenous pro-fibrotic substances than macrophages.

### Endothelial cell response to exogenous pro-fibrotic stimulus *in vitro* recapitulates known events in relevant adverse outcome pathways

To link the observed transcriptomic changes to higher-order biological effects and to contextualise them with respect to potential health outcomes, we integrated adverse outcome pathways (AOPs) in our analysis. AOPs offer a multiscale perspective connecting molecular responses to cellular, tissue, and organism-level outcomes. In brief, AOPs systematically map the progression of biological events from a molecular initiating event (MIE) at the smallest scale (e.g., receptor binding or DNA damage) through cellular, tissue, and organ-level changes organised in key events (KEs), culminating in an adverse outcome (AO) at the organism or population level. This approach connects processes across scales, enabling a comprehensive understanding of how molecular-level perturbations propagate to impact the whole system.^35^ Thus, the integration of omics data with AOPs enables a robust and multiscale analysis framework for understanding the broader implications of molecular changes. Currently, eight AOPs for PF are deposited in the AOP-Wiki repository (aopwiki.org). Although most of them are still under development and miss suggestions of prototypical stressors, neither of the MIEs of those AOPs is directly and clearly linked to endothelial cells. A recent effort to update biological information of KEs failed to annotate specific cell types to MIEs of PF, underscoring the multiplicity in PF initiation.^36^

We investigated how the transcriptomic dysregulation observed in HUVEC cells upon exposure to bleomycin and TGF-beta maps to the AOP network. We performed an enrichment analysis against the recently curated gene sets for 969 KEs of 231 AOPs using genes that were both dose-dependently and differentially expressed under bleomycin and TGF-beta, respectively.^36,37^ For bleomycin, 293 KEs part of 185 AOPs were enriched, with 49 MIEs, 206 KEs, and 57 AOs (Table S6). For TGF-beta, 124 KEs part of 125 AOPs were enriched with 10 MIEs, 88 KEs, and 32 AOs (Table S6). We explored which AOPs have the highest fraction of their KEs enriched by the two exposures. Under TGF-beta, 4 of the 8 existing PF AOPs were among the top covered AOPs, while for bleomycin only one was among the top 15 covered AOPs (Figures S5A and S5B). AOPs for TGF-beta were focused on lung and liver diseases, while for bleomycin, cardiac outcomes and various forms of cancer, glucocorticoid system/receptor activation, and male fertility were among the top enriched outcomes challenging a one-chemical one-endpoint view. Here, we focused on the early mechanism of action (MOA) of exogenous pro-fibrotic stimuli on endothelial cells and how this relates to PF as adverse outcome. Thus, we investigated the AOP subnetwork of PF and the enriched parts under bleomycin and TGF-beta. AOP:173, the only OECD-endorsed AOP for PF to date, was affected by bleomycin at multiple KEs, surprisingly not only early KEs, but also anatomical-scale KEs such as accumulation of collagen and the adverse outcome PF itself (Figure 2A). Multiple, but especially later parts of PF AOPs were fully enriched (at each timepoint) by genes altered through bleomycin exposure as prototypical exogenous pro-fibrotic stimulus, while it was striking that some KEs in the PF AOPs were systematically skipped in the enrichment including KEs such as epithelial to mesenchymal transition (EMT), increased activation of T (T) helper (h) type 2 cells, and loss of alveolar capillary membrane integrity (Table S7). This highlights that individual cell types such as endothelial cells are representing only one part in the development of a phenotype, while KEs and KE relationships are determined by intertwined networks of multiple cell types within a tissue. Although under bleomycin, the transcriptomic shift comprised a distinct signature indicating mesenchymal transition (later in the article), the KE EMT, which refers to mesenchymal transition occurring in epithelial cells, was not enriched. This underscores that extrapolations from the epithelial to the endothelial barrier are limited and both cell types come with specific responses to pro-fibrotic substances further emphasising the need of studying specific mechanisms of the endothelium to model systemically initiated PF. Besides inflammation and recruitment of immune cells, however, the KEs collagen accumulation and increased fibroblast proliferation, and mesenchymal differentiation were enriched suggesting that endothelial cells adopt mesenchymal characteristics without the involvement of fibroblasts. Further, the gene set of PF itself as AO was enriched by genes altered at any measured timepoint including 24h as extremely early timepoint with view on the development of organ fibrosis. We observed a comparable pattern in the other AOPs for PF, which naturally share many KEs with AOP:173 (Table S7). Despite the limitation of using AOPs to model chains of events using only a single cell type, we find it remarkable that under exposure to pro-fibrotic stimuli, endothelial cells beside early initiating events show multiple putatively later signatures of PF after a short duration of exposure. Interestingly, under TGF-beta, primarily later KEs of PF were consistently enriched, while early KEs related to pro-inflammatory conditions were not enriched suggesting that TGF-beta cannot be seen as a prototypical initiator of pro-fibrotic conditions, but rather a maintaining factor during the pathogenesis (Table S7).

**Figure 2:**
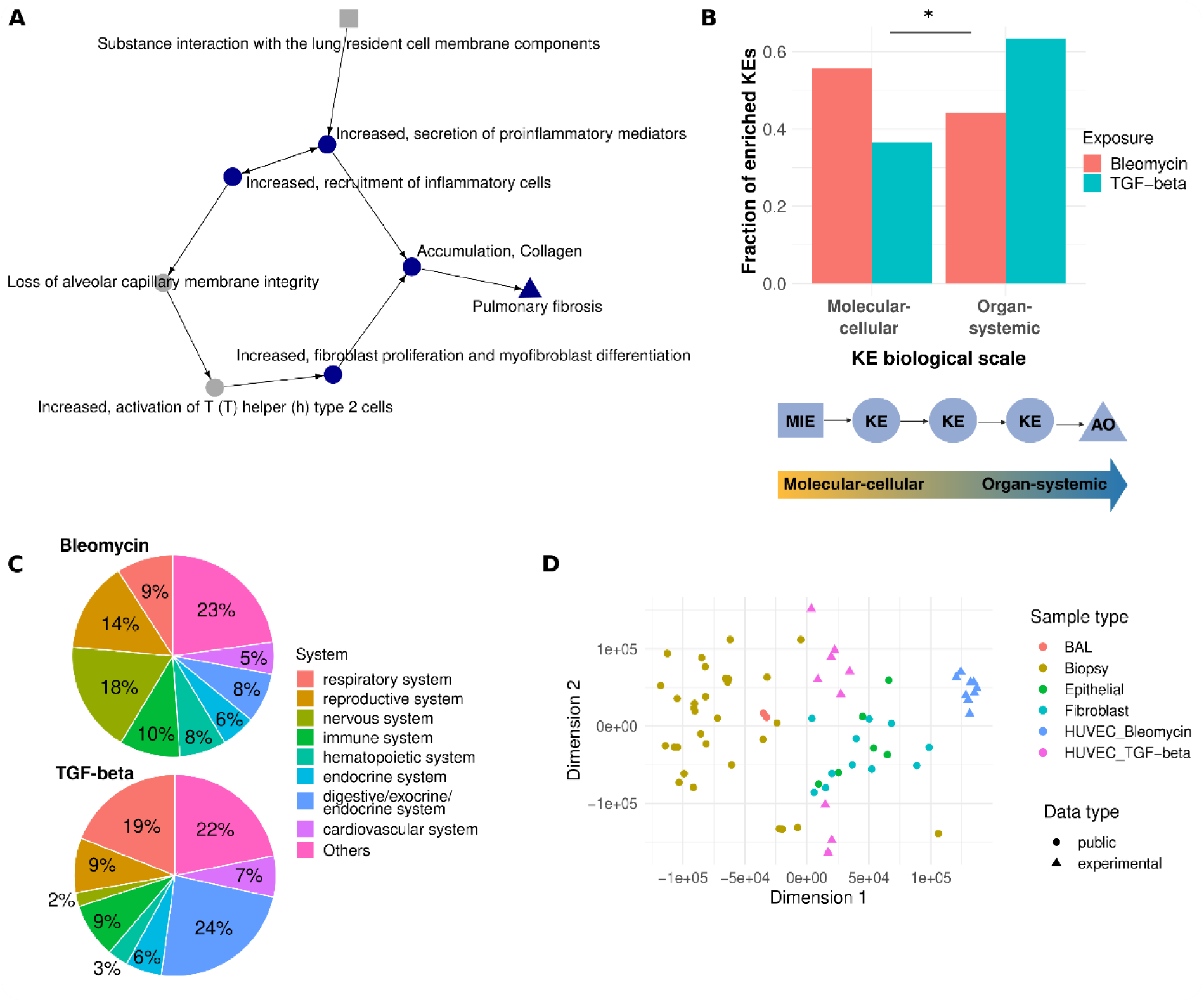
Bleomycin is a molecular initiator of fibrotic conditions. **A.** AOP:173 (Substance interaction with the pulmonary resident cell membrane components leading to pulmonary fibrosis). KEs in blue were enriched by dysregulated genes under bleomycin at each of the 3 timepoints. KEs in grey were not enriched at any timepoint under bleomycin exposure. The square indicates the MIE, the triangle the AO. **B.** The fraction of enriched KEs under Bleomycin and TGF-beta exposure that are of molecular-cellular and organ-systemic biological scale, respectively. KEs of molecular and cellular biological level were merged, presenting MIEs and early KEs of an AOP, while KEs of tissue, organ, and individual biological level were considered as of organ-systemic biological scale, presenting later KEs and AOs of an AOP. Difference between the two substances was tested by two-sided Fisher’s exact test (p-value = 2e-11). **C.** The systems of enriched KEs as recently annotated by Saarimäki *et al.* for bleomycin (top) and TGF-beta (bottom). **D.** Experimental data in comparison with public RNA-Seq data sets of PF. Sixteen public data sets and in total 51 comparisons were recently curated by Inkala *et al. (*https://doi.org/10.5281/zenodo.10692129). Fold changes of samples of PF versus healthy controls were compared with fold changes computed for HUVEC cells exposed to bleomycin and TGF-beta, respectively. The multi-dimensional scaling plot was generated from the Euclidean distance between datasets based on the ranks of 8,127 common genes according to their fold changes. BAL: bronchoalveolar lavage.

To investigate if the overall enriched KEs under bleomycin and TGF-beta exposures differ in terms of biological scale, we compared the fraction of enriched molecular and cellular KEs with the fraction of higher-level organ-systemic KEs comprising KEs on the tissue, organ, and individual level. While 56% and 37% of enriched KEs under bleomycin and TGF-beta were of molecular-cellular biological scale, 44% and 63% were of organ-systemic scale, respectively, suggesting that bleomycin as prototypical exogenous pro-fibrotic substance is primarily a molecular initiator, whereas TGF-beta is a downstream regulator of cellular networks and thus more actionable in higher organ or tissue (dys)functions with intertwined actions of multiple cell types (Figure 2B). The most covered body systems under TGF-beta were digestive and respiratory system, while for bleomycin it was nervous system and reproductive system, likely given the known genotoxic role of bleomycin (Figure 2C). Interestingly, the reproductive and nervous system are two systems that have a specialised endothelium preventing an uncontrolled entrance of molecules from the circulatory system, while liver and lungs are, by function, very accessible.

Beside using the AOP framework to place molecular findings on a broader and phenotype-related level, we contextualised our *in vitro* observations with real-world PF scenarios. Therefore, we made use of a recently curated data repository of clinical PF transcriptomic datasets (https://doi.org/10.5281/zenodo.10692129). We used precomputed fold changes of those public datasets resembling transcriptomic changes in individuals with PF as compared to healthy controls and compare them with our observed transcriptomic changes in *in vitro* exposed endothelial cells as compared to unexposed cells (Figure 2D). Whole-tissue samples such as biopsies and bronchoalveolar lavages showed differences to isolated cell populations from clinical samples such as epithelial cells and fibroblasts. Though our data were derived from isolated cell cultures that have never been part of a cellular network in a fibrotic organ, HUVEC cells exposed to TGF-beta showed comparable transcriptomic shifts to epithelial cells and fibroblasts derived from lung fibrosis. Since constant and exaggerated attempts of tissue repair is a hallmark of fibrosis, TGF-beta is one of the key maintainers of organ fibrosis.^4^ This might explain why short-term exposure of endothelial cells to TGF-beta establishes a transcriptomic profile similar to other cells from a fibrotic environment. Bleomycin exposure overall resulted in a different transcriptomic profile underscoring the importance of differentiating between MOAs in established fibrotic conditions and early pro-fibrotic stimuli by exogenous compounds.

Altogether, TGF-beta-exposed endothelial cells showed transcriptomic profiles closely related to established fibrotic states as demonstrated by clinical data of PF and higher level KEs in AOPs of PF. In contrast, bleomycin-exposed cells presented a distinct signature, highlighting its role as a potent and hazardous molecular stressor, but already after short time of exposure, signatures of developing fibrosis can be observed.

### Bleomycin shifts the overall transcriptomic fingerprint of HUVEC cells caused by damage-associated molecular patterns

The described direct MOA of bleomycin, while not inherently linked to its pro-fibrotic effects, is (1) lipid peroxidation of membrane fatty acids and (2) binding DNA leading to DNA breaks.^38^ Exploring the transcriptome-wide dysregulation under bleomycin exposure indeed revealed downregulation of DNA replication and cell cycle genes at every timepoint and concentration of bleomycin, as well as a disturbance of DNA repair mechanisms at the first day of exposure which normalised over time (Figure 3A, Table S8). In accordance, the p53 pathway was upregulated at all measured timepoints, commonly observed upon DNA damage to halt cell cycle and in case DNA repair fails to induce apoptosis.^39^ Further, MTORC1 signalling was downregulated at 24h of bleomycin exposure indicating a compromised nucleotide and lipid biogenesis, which however recovered at later timepoints (Figure S6, Table S8).^40^ In contrast to the downregulated fingerprint, the profile of upregulated genes under bleomycin clearly indicated an inflammatory response involving TNF alpha and cytokine signalling (Figure 3A, Table S8). Further, upregulated genes were enriched in pathways related to ECM receptor interaction and adhesion molecules indicating a shift in ECM organisation and cell junctions. Enriched pathways such as myogenesis and EMT, despite referring to signatures described in other cell types than endothelial cells, suggest a shift in the transcriptome of endothelial cells towards a mesenchymal-like cell type. After 48h, hallmark pathways such as interferon alpha and gamma response were upregulated, likely representing an internal cellular mechanism for immune surveillance of DNA damage (Figure S6, Table S8).^41^

**Figure 3:**
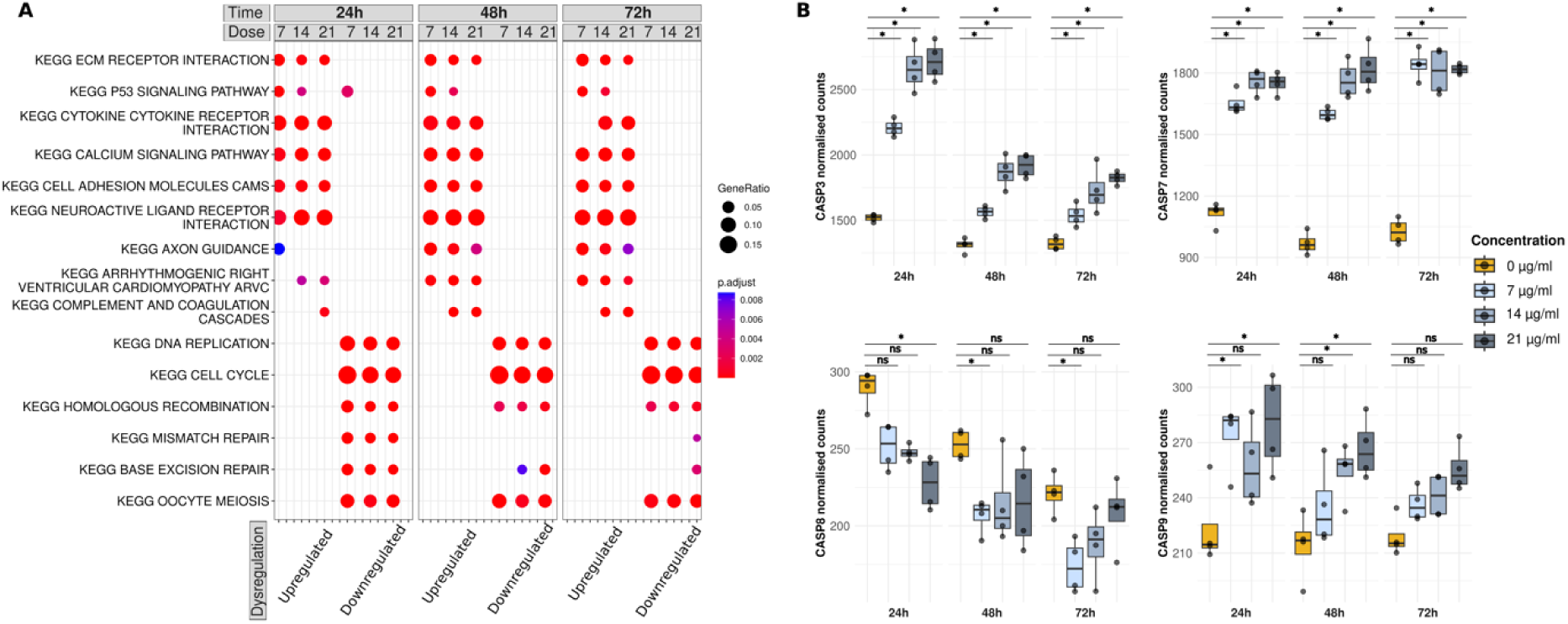
Transcriptomic changes under bleomycin exposure. **A**. Functional analysis of up-and downregulated genes under bleomycin exposure. Overrepresentation test with dysregulated genes (absolute LFC > 1, adjusted p-value < 0.01) against the gene sets of KEGG pathways. The dot size indicates the gene ratio *i.e.*, the ratio of identified genes to pathway genes. The colour code indicates the Benjamini-Hochberg-adjusted p-value of the Fisher’s exact test. The top five enriched pathways are plotted, and already existing pathways are filled even if they are not among the top five pathways. **B.** Expression of apoptosis marker genes for the different experimental conditions of bleomycin exposure. Plotted are the size-factor normalised RNA-Seq counts.

Despite the use of subtoxic doses, we observed an induction of apoptotic effectors on the transcriptome level. Caspase 3, 7, and 9 (*CASP3*, *7*, *9*) encoding genes involved in the activation of the intrinsic pathway of apoptosis were upregulated at all concentrations and timepoints of bleomycin as compared to unexposed cells, whereas *CASP8* involved in the extrinsic pathway of apoptosis became even downregulated under bleomycin (Figure 3B).^42^ The intrinsic pathway of apoptosis is initiated when cells encounter intracellular stress such as in response to DNA damage due to bleomycin.^43^ Further, it was shown that bleomycin accumulates in mitochondria, damages mitochondrial DNA and affects mitochondrial integrity further promoting the intrinsic pathway of apoptosis.^38,44^ Both nuclear and mitochondrial DNA are sensed as damage-associated molecular patterns (DAMPs) and promote wider downstream effects on the cell itself as well as neighbouring cells such as the induction of controlled or uncontrolled cells death.^41^ Nuclear DAMP receptors sensing nuclear DNA such as *CGAS* and *AIM2* were upregulated at later timepoints and higher concentrations of bleomycin only, while *RIGI* encoding a cytosolic DAMP receptor (RIG-I) for mitochondrial DNA was upregulated already at the lowest concentration and 24h of exposure suggesting that mitochondrial DNA damage might even precede nuclear DNA damage (Figure S7).^44^

Since bleomycin initiates decomposition of membrane fatty acids, a destabilised membrane might further lead to cell leakage and thus resulting in intercellular DAMP sensing by membrane receptors of neighbouring cells such as TLR2 and TLR4, both of whose transcription was impelled immediately after bleomycin exposure (Figure S7).^41^ A direct downstream effect of TLR2/4 signalling is the production of pro-inflammatory cytokines as we observed it on the transcriptomic level (Figure 3A). Despite clear induction of apoptotic machinery, including early downregulation of *BCL2* encoding a suppressor protein of apoptosis, the normalisation of *BCL2* expression levels in cells exposed to higher doses of bleomycin from 48h on indicates an adaptive response to resist further apoptotic signals enabling cell survival (Figure S7).^45^

In summary, bleomycin-induced DNA damage and lipid peroxidation is a source for DAMPs, triggering apoptosis, which subsequently generates additional DAMPs that amplifies the inflammatory response of endothelial cells. This chain of events offers a possible link of bleomycin’s direct MOA to its observed pro-fibrotic effects.

### PF associated gene signatures can be observed in HUVEC cells exposed to exogenous pro-fibrotic stimulus, but not TGF-beta

To understand the mechanistic events that eventually promote a pro-fibrotic environment starting from a dysregulation of endothelium, we investigated reviewed key cellular processes of PF propagation related to endothelial cells by transcriptomic signatures and cytokine secretion (Figure 4A).

**Figure 4:**
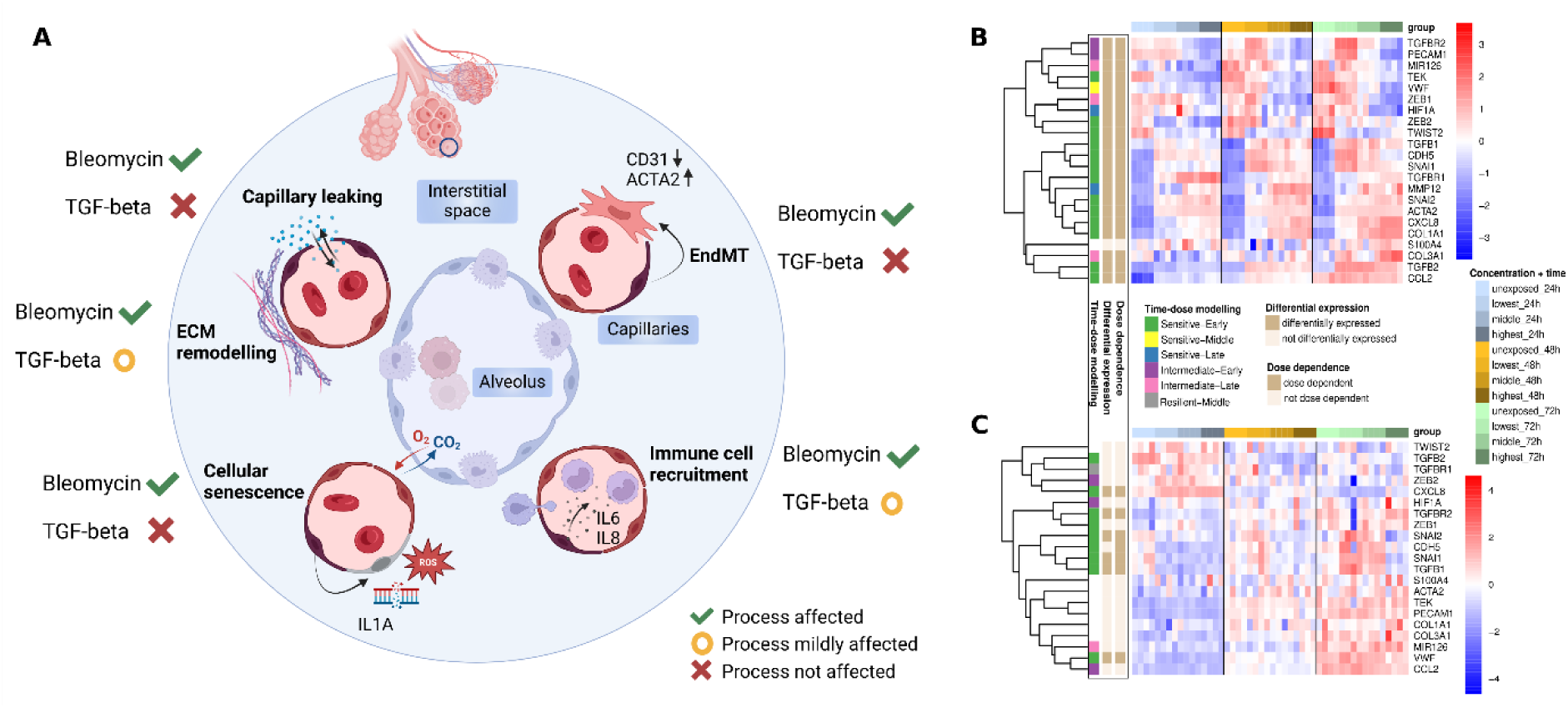
Fibrosis-associated cellular processes in endothelial cells. **A.** Cross-section of an alveolus and the surrounding capillaries highlighting key endothelial cell processes associated with PF. Processes are annotated as follows: a green checkmark indicates that RNA-Seq data showed the process was affected by bleomycin and TGF-beta exposure, respectively; a red cross indicates the process was not affected; and a yellow circle indicates that the process was only mildly or transiently affected. **B.** Marker genes of EndMT under bleomycin exposure. The colour annotation of columns indicates the experimental condition, the row annotations refer to the label of the time-dose modelling, and the results of the differential expression and dose-dependent analysis. The heatmap colour code represents row-wise z-scores based on vst-transformed counts. **C.** Marker genes of EndMT under TGF-beta exposure. Annotations and colour codes are the same as described in B.

EndMT is a process where endothelial cells undergo the transition towards a mesenchymal phenotype characterised by increased expression of mesenchymal markers such as *ACTA2*, decreased expression of vasculature-specific genes such as *PECAM1* (encoding CD31), and loss of barrier integrity.^27,46^ Under bleomycin, EndMT markers were overall differentially expressed in HUVEC cells, and all of them were dose-dependently altered (Figure 4B), while time-dose modelling indicated that most genes responded early sensitive *i.e.*, they are changing early (24h) and at low subtoxic doses already (Table S9). Beside commonly investigated markers for EndMT, we observed an induction of *MYOCD*, a gene specific for smooth muscle cells (Figure S8).^29^ *MYOCD* was neither detectable in unexposed HUVEC cells nor when they were exposed to TGF-beta. TGF-beta signalling, central to EndMT and fibrotic conditions, was induced under bleomycin as shown by upregulation of *TGFB1*, *TGFB1R*, and *SMAD3*, but downregulation of TGF-beta-signalling inhibitory *SMAD6* (Figure S8).^47,48^ In contrast to bleomycin, exposure of the cells to exogenous TGF-beta 1 itself led to an initial suppression of TGF-beta signalling with later recovery likely reflecting a negative feedback loop (Figure S8). Importantly, EndMT marker genes including landmark mesenchymal transition markers such as *ACTA2* and *PECAM1* did not indicate a progression of HUVEC cells to a more mesenchymal-like cell type under TGF-beta exposure within 24 to 72 hours (Figure 4C).

In PF, endothelial dysfunction and capillary leakage are linked to disrupted cell-cell junctions, early markers of EndMT.^48^ Bleomycin exposure caused dose-dependent dysregulation of cell-junction related genes, including suppression of *CLDN5*, a key tight junction component, consistent with observations in bleomycin-PF mouse models (Figure S9).^21,49^ In contrast, TGF-beta exposure did not affect the expression of cell-cell junction components including *CLDN5*, contradicting observations of Ohta *et al*.^21^As shown above, both bleomycin and TGF-beta exposure altered genes enriched in ECM related processes and functions. Differential expression and altered protein secretion of *PDGFB* and *MMP7* further indicated that cells underwent ECM remodelling processes upon exposure with bleomycin, a hallmark of fibrosis development and concomitant mesenchymal transition of cells (Figure S10).^4^ Besides, PDGF is a chemoattractant for fibroblasts.^50,51^

Cellular senescence occurs when apoptotic signals are insufficient, and environmental conditions inhibit active cell cycle and metabolism.^52^ Using the SenMayo gene set, we found widespread dose-dependent induction of senescence-associated genes under bleomycin exposure, whereas TGF-beta had only minor effects (Figure S11).^53^ In endothelial cells, interleukin-1 alpha was shown to indicate cellular senescence, and we observed both a dose-dependent upregulation of *IL1A* on the transcript level and a dose-dependent increase of secreted IL-1 alpha under bleomycin, while IL-1 beta secretion was undetectable (Figure S12).^54^ Interestingly, while comparable concentrations of TGF-beta were shown to induce senescence in human bronchial epithelial cells, endothelial cells appeared more resistant to such effects of TGF-beta.^55^

Key chemoattractants for monocyte recruitment such as IL-6 and IL-8 showed an early dose-dependent upregulated expression and increased secretion under bleomycin exposure (Figure S13).^56,57^ TGF-beta, however, had only transient and dose-independent effects on IL-6 and IL-8 transcript levels and did not influence their secretion at the 3 measured timepoints, further indicating self-limited effects of TGF-beta on endothelial cells. We acknowledge that especially for TGF-beta exposure, 24h might be already too late to detect early MOAs given that we observed a reversion of signals already at 48h.

Altogether, our findings demonstrate that bleomycin as prototypical exogenous pro-fibrotic substance induces EndMT- and senescence-associated gene profiles, accompanied by signatures of disturbed cell-cell junctions. An increased secretion of IL-6 and IL-8 under bleomycin suggests the ability of endothelial cells to effectively recruit monocytes from the blood stream early after contact with this exogenous pro-fibrotic compound. While TGF-beta altered expression levels of genes related to the ECM remodelling machinery, its effects on endothelial cells appear minor and time-limited.

## Conclusion

Our findings highlight the exceptional sensitivity of endothelial cells to subtoxic doses of bleomycin, a prototypical exogenous pro-fibrotic compound, with pronounced transcriptomic alterations observed shortly after exposure. Bleomycin emerged as an early initiator of pro-fibrotic signals without the contribution of other cell types like fibroblasts and monocytes. Endothelial cells displayed signatures characteristic of established organ fibrosis, including profiles of EndMT, cellular senescence, and immune cell recruitment. In contrast, TGF-beta exposure resulted in minimal and self-limited transcriptomic effects, suggesting its pro-fibrotic activity may require a cellular network involving fibroblasts or prior stimulation by exogenous factors. Given the limitations of studying the transcriptome, we explored multiple timepoints and doses of exposure and complemented this with studying secretion of candidate proteins. Nonetheless, transcriptomic snapshots can only be used to interpolate a dynamic expression landscape and derive a MOA. Bleomycin-induced DNA damage, particularly in mitochondria, initiates intrinsic apoptotic pathways, with damaged DNA acting as a source of DAMPs that drive inflammation and mesenchymal transitions. Three days of subtoxic bleomycin exposure provide a blueprint for how endothelial cells reprogram their transcriptional activity toward pro-fibrotic and pro-inflammatory processes, laying the foundation for extensive downstream effects within the tissue context. While endothelial cells may not be the primary drivers of fibrosis, their role in its initiation is likely underestimated compared to epithelial cells and fibroblasts. Our study highlights the critical early initiating events triggered by an exogenous pro-fibrotic substance upon contact with endothelial cells, underscoring their potential as potent initiators of pro-fibrotic mechanisms.

## Methods

### Cell culture

Human umbilical cord blood vein cells (HUVEC) were ordered from ATCC. Cells were cultured in Endothelial Cell Growth Medium-2 (EGM-2, Promocell #C-22011). Prior to the experiment, cells were expanded in T-75 flasks for 1-2 passages. For the experiments, passages 6-8 were used and cells were plated on 12-well (for transcriptomics) or 96-well (for viability) plates with a density of 40 000 cells/cm^2^. Cells were settled for 24h prior to experimentation.

### Exposures

The cells were exposed to bleomycin (Sigma-Aldrich, #B7216) in 96-well plates for conducting a viability assay using concentrations of 0 to 520 µg/ml with continuous exposure. Thereafter, subtoxic concentrations of 7, 14, and 21 µg/ml bleomycin were used in 12-well plate format for further analyses corresponding to IC5, IC10 and IC20 (Figure S14).

TGF-beta (Sigma-Aldrich #T7039) was used in concentration of 0 to 100 ng/ml for conducting a viability assay. TGF-beta was not compromising the viability of the cells. Therefore, concentrations of 0, 5, 10, 20 ng/ml TGF-beta were used in 12-well plate format according to concentrations found in the literature and which mimic physiologically relevant concentrations given that the cells were exposed from their apical site.^33,58–60^

### Viability assay

Real time Glo MT Cell Viability Assay (Promega, G9713) was used to determine the viability of the cells. The assay was mixed into cell culture medium and added to the cells together with exposure. Luminescence was measured after 72h incubation in +37°C, and 5% CO_2_. Luminescence was measured using a Spark microplate reader (Tecan).

### Immunoassay against cytokines and peptidases

Procartaplex immunoassay was designed for potential secreted factors important to pro-fibrotic processes. Selected cytokines and peptidases were: CXCL11, FGF-2, IFN gamma, IL-1 alpha, IL-1 beta, IL-10, IL-13, IL-17A, IL-18, IL-6, IL-8, IP-10, MCP-1, MIG, MIP-1 alpha, MMP-1, MMP-7, MMP-9, PDGF-BB, SDF-1 alpha, TNF alpha, and VEGF-R2. Immunoassay was run from supernatant collected from bleomycin or TGF-beta exposed cells at 24h, 48h, and 72h, respectively. The assay was read using a Luminex 200 (BioRad). Statistical testing was performed by ANOVA and pairwise t tests. Differential secretion was defined by a false discovery rate (as defined by Benjamini-Yekutieli) of <5%. Prism (v10.3.1, Graphpad) was used for both testing and visualisation.

### Dose-dependent analysis of cytokines and peptidases

Dose-dependent cytokine and proteinase secretion under bleomycin exposure was assessed using the BMDx software.^61^ Only proteins with 4 replicates across all conditions (4 doses and 3 timepoints) were analysed. The relative intensities from the immunoassay were log-transformed using the formula log2 (x+1) to normalise the data and reduce skewness. A list of dose-response models up to the third degree was fitted to the data of 4 doses per timepoint including linear, polynomial, and nonlinear types (e.g., Weibull, exponential). Models were filtered using an R-squared (R²) threshold of 0.6, which is a suitable trade-off between variance of the data and complexity of the model. For each protein, a representative optimal model was selected as the one with the minimum Akaike Information Criterion value, since it provides a good compromise between goodness of fit and model complexity. Only proteins with at least one model after filtering are deemed dose-dependent.

### RNA extraction

RNA was extracted from cells in 12-well plates after 24h, 48h, and 72h of exposure using the RNeasy Mini kit (Qiagen). Four parallel samples were collected per experimental group. Prior to cell lysis, supernatant was collected for cytokine secretion assay. DNase 1 (Thermo Scientific, EN0521) treatment was used prior to downstream analyses.

### RNA sequencing (RNA-Seq)

RNA-Seq was performed in two batches namely the two exposure experiments of bleomycin and TGF-beta, respectively. Prior to library preparation, the quality of the samples was determined by gel electrophoresis using Bioanalyzer 2100 (Agilent Technologies, USA). Library preparation and RNA sequencing was conducted by the company Novogene using the protocols described earlier.^34^ Libraries were sequenced using the Illumina NovaSeq X platform, according to effective library concentrations and data amounts using the paired-end 150 strategy (PE150). Library preparation and sequencing was repeated for 3 samples of the TGF-beta experiment (C_91, C_96, and C_173) due to exceptionally low CG content in the first sequencing datasets (Table S10). On average, 24 and 28 million read pairs were generated per sample in the bleomycin and TGF-beta experiment, respectively (Table S10).

### RNA-Seq data processing

Sequencing data quality was assessed using FastQC (v0.12.1, https://www.babraham.ac.uk) and summarised with MultiQC (v1.7).^62^ Raw RNA-Seq reads were trimmed for low-quality bases (Phred quality score < 20) and Illumina sequencing adapters using Cutadapt (v4.4).^63^ Reads shorter than 60 bases post-trimming were discarded. Alignment of the trimmed sequencing reads to the Genome Reference Consortium Human Build (GRCh38) Ensembl version 110 was performed using HISAT2 (v2.2.1) with default settings.^64^ Bam files were filtered for uniquely mapped reads (as flagged by “NH:i:1”) and sorted by coordinates using Samtools (v1.18).^65^ To quantify expression per gene, read summarisation was conducted with the featureCounts function from the Rsubread package (v2.16.1) against the Ensembl GTF file version 110.^66^

### RNA-Seq analysis

Count matrices were handled and processed in R using DESeq2 (v1.36.0) functions.^67^ Counts normalised for the effective RNA-Seq library size (SizeFactors as computed by DESeq2) were used to illustrate gene expression heights in gene plots using the plotCounts function. For other downstream analyses, unless explicitly mentioned, counts were normalised and transformed using the variance stabilising transformation (vst) function from DESeq2 after filtering for genes with at least 10 counts in a minimum of 4 out of 48 samples (due to replicates of 4). Vst was performed in a condition-aware manner (blind parameter set to FALSE). Principal component analysis was performed to assess overall data quality and global data structure.

Differential expression analysis was performed using the parametric Wald test and subsequent independent filtering of the results as implemented in DESeq2.^67^ For each of the two experiments (bleomycin and TGF-beta exposure), pairwise comparisons were conducted between samples of exposed cells (combination of timepoint and concentration) versus unexposed controls. This approach allowed for the identification of gene expression changes specific to each experimental condition. Differentially expressed genes were generally defined by a false discovery rate (as defined by Benjamini-Hochberg) of <1% without any further filter regarding fold change. Since expression changes under bleomycin were vast (in total 18,531 differentially expressed genes), a fold change filter was used in two cases of enrichment tests as described further below.

Dose-dependent gene expression analysis was conducted with 4 replicates at 4 doses for each timepoint using the BMDx tool.^61^ Since 4 doses of exposure were tested, statistical models up to the third degree were fitted to the vst-transformed counts including linear, Hill, power, polynomial, log-logistic, and Weibull models. An R-squared (R²) threshold of 0.6 was applied ensuring a strong and reliable dose-response relationship. For each gene, the model with the lowest Akaike Information Criterion was selected as the best fit.

AOP enrichment analysis using the gene-key event (KE) annotation as published in Saarimäki *et al.* was performed as described recently.^36,37^ The intersection of dose-dependent genes (at least at one timepoint) and differentially expressed genes (at least at one timepoint and one dose) under bleomycin and TGF-beta, respectively, was used for KE enrichment analysis. Given the high degree of dysregulation under bleomycin, an absolute LFC threshold of >1 corresponding to doubled or halved expression [log_2_(2) = 1, log_2_(0.5) = -1] was applied as further criterion to define differentially expressed genes under bleomycin (yielding 6,833 genes in total as input for the enrichment analysis). Since effects of TGF-beta on HUVEC cells were overall small, we analysed the data deliberately without applying a LFC filter (yielding 281 genes in total as input for the enrichment analysis). The fraction of KEs enriched in each AOP was calculated and the top 15 enriched AOPs were illustrated. The biological annotation of AOPs and KEs as published in Saarimäki *et al.* was used to define KEs of lower- and higher-level biological scale.^36^ In detail, KEs of molecular and cellular biological level were merged, presenting MIEs and early KEs of an AOP, while KEs of tissue, organ, and individual biological level were considered as of organ-systemic biological scale, presenting later KEs and AOs of an AOP. Network visualisation of AOP subnetworks was performed with the igraph R package (doi:10.5281/zenodo.7682609).

Time-dose modelling was conducted using the TinderMIX R package.^68^ Pairwise LFC for each gene, computed as the difference between the vst counts of each pair of exposed and unexposed samples per timepoint were used to fit a linear and 2nd- and 3rd-order polynomial models. Model selection was performed using a nested analysis of variance (ANOVA) and the best fitting model was used to predict an activity map (contour plot) for each gene. The dynamic dose-responsive area was identified under the condition that a 10% increase or decrease with respect to controls was reached (activity threshold for the benchmark-dose modelling). To label each gene with an activation label specifying its point-of-departure, the lowest dose and earliest timepoint of activation was identified, and the gene was labelled as “sensitive”, “intermediate”, or “resilient” regarding the dose, and “early”, “middle” or “late” regarding the time.

A recently curated data repository of clinical PF transcriptomic datasets was used to contextualise the observed transcriptomic changes under the exposure with changes observed in real-life PF specimens (https://doi.org/10.5281/zenodo.10692129). Precomputed fold changes of those public datasets resembling transcriptomic changes in individuals with PF as compared to healthy controls were compared with the observed transcriptomic changes in *in vitro* exposed endothelial cells as compared to unexposed cells. In detail, LFCs for each gene as computed by DESeq2 were retrieved for each comparison of the clinical datasets (n=51 comparisons) and of the *in vitro* datasets *i.e.*, 9 comparisons for bleomycin and TGF-beta, respectively (3 timepoints and 3 doses). The 8,127 genes the 69 comparisons had in common were ranked according to the LFC in each comparison. Based on the gene ranks, Euclidean distance was computed between the comparisons and visualised in a multi-dimensional scaling plot.

Functional enrichment analysis of differentially expressed genes under bleomycin was conducted using the clusterProfiler R package (v4.4.4) with gene sets from Gene Ontology (GO) and the Molecular Signatures database (MSigDb) including the KEGG pathway and Hallmark pathway sets.^69–75^ Overrepresentation tests were performed for each set of differentially expressed genes for an experimental group and an absolute LFC threshold of >1. For the functional enrichment analysis of dysregulated genes that bleomycin and TGF-beta exposure had in common, differentially expressed genes were not filtered by fold change for both exposures, as TGF-beta had overall small and narrow effects on the cells and a fold change filter did not leave enough genes for functional analysis. Overrepresentation test was performed with overlapping differentially expressed genes at each timepoint under the two exposures against the GO gene sets using clusterProfiler.^69,70,74^

Candidate gene sets reflecting signatures of key endothelial processes associated with PF were determined based on literature in the field.^27,46,53,76–78^ For the exploration of changes under TGF-beta exposure, genes that were significantly dose-dependently altered under TGF-beta at all timepoint were uploaded to the STRING database in the “multiple protein” mode to derive protein-protein-interactions from the STRING interactome.^79^ RNA-Seq data analysis was performed using R (v4.2.1). Visualisations were performed with R, Biorender, and Inkscape.

## Abbreviations

AO: Adverse outcome
AOP: Adverse outcome pathway
DAMP: Damage associated molecular pattern
ECM: Extracellular matrix
EMT: Epithelial to mesenchymal transition
EndMT: Endothelial to mesenchymal transition
HUVEC: Human umbilical vein endothelial cells
KE: Key event
LFC: Log2 fold change
MIE: Molecular initiating event
MOA: Mechanism of action
PF: Pulmonary fibrosis
vst: Variance stabilising transformation

## Declaration of generative AI and AI-assisted technologies in the writing process

During the preparation of this work the authors used ChatGPT to improve the readability of single sentences and to shorten single paragraphs with limited word count allowed. After using this tool, the authors reviewed and edited the content as needed and take full responsibility for the content of the published article.

## Author contributions

L.Mö.: Conceptualisation, Methodology, Software, Validation, Formal Analysis, Investigation, Data Curation, Writing – Original Draft, Writing – Review & Editing, Visualization; L.Y.: Methodology, Validation, Formal Analysis, Investigation, Writing – Original Draft, Writing – Review & Editing, Visualization; L.Ma.: Methodology, Formal Analysis, Investigation, Writing – Review & Editing; G.M.: Methodology , Investigation, Writing – Review & Editing, Visualization; K.K.: Investigation, Writing – Review & Editing; N.D.: Investigation, Writing – Review & Editing; A.S.: Methodology, Writing – Review & Editing , Supervision; D.G.: Conceptualisation, Resources, Writing – Review & Editing, Supervision, Project Administration, Funding Acquisition

